# Quercetin Suppresses TNBC Cell by Targeting ORM2

**DOI:** 10.1101/2023.06.20.545736

**Authors:** Zhijun Chen

## Abstract

**Background:** Triple-negative breast cancer (TNBC) is known for its aggressive nature, and Quercetin (QUE) has shown potential anti-cancer effects.

**Methods:** We determined the IC50 of QUE for inhibiting cell viability in multiple TNBC, non-TNBC, and normal breast cell lines. We compared the expression of ORM2 in TNBC clinical samples and normal tissues. Additionally, we measured ORM2 expression in TNBC and normal breast cell lines. We determined the IC50 of QUE for inhibiting cell viability after ORM2 knockdown. An orthotopic implantation mice model was used to evaluate the treatment effect of QUE. We also conducted molecular docking and amino acid exchange validation to model the binding of QUE to ORM2. Furthermore, we performed a protein-protein interaction network analysis and GO enrichment analysis of differentially expressed genes associated with ORM2 in TNBC.

**Results:** QUE inhibited the viability of both TNBC and non-TNBC cell lines, but it was specifically associated with worse survival in TNBC patients. We observed higher expression of ORM2 in breast cancer cells compared to normal breast cells. Knockdown of ORM2 reduced the viability of TNBC cells. Treatment with QUE inhibited ORM2 expression and decreased viability in TNBC cells. In the animal model, QUE improved survival and downregulated ORM2 expression in tumors. Enrichment analysis provided insights into the potential functions of ORM2.

**Conclusion:** Our findings indicate that QUE directly inhibits TNBC cell viability through its interaction with ORM2. These results contribute to our understanding of the anti-cancer mechanisms of QUE in TNBC and highlight ORM2 as a potential therapeutic target.

## Introduction

Cancer ranks as a leading cause of death and a critical challenge for public health in the world [1]. Breast cancer is one of the most frequent cancer types diagnosed, resulting in 2,261,419 new cases and 684,996 death last year Globally[2]. According to American Cancer Society statistical data, breast cancer is the most privileged cancer type and the second leading cause of cancer-related death among women in the United States[3]. Breast cancer is a very heterogeneous cancer type[4]. Breast cancer can be categorized into three major subtypes based on immunohistochemical expression levels of estrogen receptors, progesterone receptors, and HER2: hormone receptor-positive, human epidermal growth factor receptor 2 (HER2)-enriched, and triple-negative breast cancer.

Triple-negative breast cancer (TNBC) is a highly aggressive subtype of breast cancer associated with a poor prognosis. It is characterized by the absence of estrogen receptor and progesterone receptor expression, along with overexpression of the human epidermal growth factor receptor 2 (HER2). TNBC accounts for approximately 10-15% of all breast cancer cases. [3] Most TNBC results in high-grade invasive ductal carcinoma with a higher rate of early recurrences and distant metastases[5]. The hormone has been suggested to be critical for some cancer types [6-8]. In recent years, overall breast cancer treatment has been improved significantly because of the advancement in hormone therapy (for hormone receptor-positive breast cancer) and target therapy (for human epidermal growth factor-enriched breast cancer) [9-12]. Unfortunately, so far, clinical outcomes of TNBC therapy remain unsatisfactory. The overall survival for metastatic TNBC patients is less than two years, much worse than the survival of the other breast cancer types [13]. This fact emphasizes the compelling need to explore more effective therapeutic approaches for TNBC patients.

Quercetin (QUE; 3,5,7,3′,4′-pentahydroxy flavone) has demonstrated chemopreventive effects in both in vitro and in vivo studies, particularly in the treatment of cancer. Recent investigations have revealed that QUE can induce cancer cell death through lysosome activation mediated by the transcription factor EB and reactive oxygen species-induced ferroptosis, independently of p53. Furthermore, QUE has been found to inhibit prostate cancer by suppressing cell survival and inhibiting anti-apoptotic pathways. In the context of breast cancer, QUE has shown inhibitory effects on the proliferation of various breast cancer cell lines [14]. The pro-oxidant ability of QUE, as well as the potential effect of QUE on apoptosis, was suggested to account for the prevention effects on breast cancer cells[15]. In the clinical treatment of TNBC, QUE was used as a synergistic anticancer reagent with curcumin [16]. In TNBC patients, QUE was found to impair HuR-driven progression and migration of cancer cells [17]. Another study also suggested that QUE can affect TNBC cell migration, which reported that this action was mediated by the β-catenin signaling pathway [18]. However, to date, most of the targets of QUE have not been identified. This prevents the understanding of the pharmacological effects of QUE and impedes the clinical application of QUE for TNBC treatment.

Overall, in this study, we identified a novel pharmacological target of QUE on TNBC, Orosomucoid 2 (ORM2), based on bioinformatics data screening and validated it with experimental evidence. ORM2 gene encodes a key acute-phase plasma protein. Because of its increase due to acute inflammation, this protein is classified as an acute-phase reactant[19]. However, the specific function of this protein has not yet been determined. Our study is conducive to a better understanding of QUE effects on TNBC.

## Methods and materials

### Cell lines and Cell culture

TNBC cell lines MDA-MB-231, MDA-MB-468, MDA-MB-436, BT-20, and BT-549, HER-enrich breast cancer cell line MDA-MB-435, ER-positive breast cancer cell line MCF-7, and the two non-tumorigenic epithelial breast cell lines MCF-10A and MCF-12A were purchased from the ATCC. DMEM with 10 mM HEPES, 10% FBS, and 2% pen/strep were used to culture these cells. A 37°C, 5% CO2 humidified incubator was used to culture the cells.

### Reagent and antibody

Quercetin was purchased from Sigma-Aldrich. The quercetin stock solution was prepared in methanol at 1mg/ml and sonication for 10 minutes. The MTT and trypan blue were purchased from Abcam. Human ORM2 Antibody (H00005005-B01P) and GAPDH antibody (ab8245) were used for western blotting. ORM2-Specific Monoclonal antibody (66217-1-Ig) was used for immunohistochemistry stainings of both human and mice tissue. Secondary antibodies conjugated with HRP were from Sigma-Aldrich.

### Transfection

Plasmid transfection[20] was conducted as a previous paper. ORM2 knockdown and overexpression were achieved by transfecting the ORM2 shRNA plasmid or ORM2 expression plasmid into cells. Two distinct sequences of siRNAs were used to knock down the expression of endogenous ORM2 in cells. The siRNAs sequences were cloned to the ORM2 shRNA silencing Adenovirus plasmids (Ad-h-ORM2-shRNA). The shRNA plasmid and scrambled shRNA Control plasmids were designed and constructed by the VECTOR BIOLAB (Malvern, PA, USA). The ORM2 (WT or mutated) expression plasmids (pCMV6-Entry-ORM2) and expression control plasmids were purchased from the Origene. Lipofectamine® 2000 was used to conduct the transfection experiments following the standard protocol.

### Viability assay

MTT assay[21], apoptosis ELISA[22], and cell counting[23] were conducted as previously described. Preincubate cells at 1 × 10^6^ cells/ml in culture medium at 37°C and 5% CO2. Seed cells at 5×10^4^ cells/well in a 100 μl culture medium containing testing agents into 96 wells flat bottom microplates. Incubate cell cultures for 24 h, then add 10 μl of the MTT labelling reagent (final concentration 0.5 mg/ml) to each well. Incubate the microplate for 4 h at 37°C, 5% CO2. Add 100 μl of the Solubilization solution DMSO into each well. Check for complete solubilization of the purple formazan crystals and measure the absorbance of the samples using a microplate (ELISA) reader. The wavelength to measure the absorbance of the formazan product was 550nm. CYCS/Cytochrome c ELISA Kit (LS-F11267) from LifeSpan BioSciences was used to determine the apoptosis of cells following the instructions of the kit. Cells were resuspended and cell numbers were counted using a hemacytometer with staining of trypan blue.

### Western blotting assay

Western blotting assay was conducted as a previous paper**[24]**. Samples were homogenized in reducing sample buffer (80 mM Tris–HCl [pH 6.8], 0.1 M dithiothreitol, 2% sodium dodecyl sulfate, and 10% glycerol) and Halt phosphatase and protease inhibitor cocktail (Thermo Scientific) using a Dounce homogenizer. An equal amount of total protein for each sample was loaded on sodium dodecyl sulphate-PAAG followed by transfer to a nitrocellulose membrane. Specific signals were developed using ChemiGlow chemiluminescent substrate for HRP (Protein Simple). Images of the blots were acquired using FluorChem FC2 Imager (Alpha Innotech). Quantitative analysis was performed using ImageJ software.

### Immunohistochemistry stainings

Immunohistochemistry stainings were conducted as a previous paper [25]. Tissue microarray (TMA) blocks were made with 50 cases of TNBC with adjacent normal tissues or made with animal tumor samples using 3 mm diameter cores. Immunohistochemistry with antibodies was performed on the BondMAX automated immunohistochemistry staining platform using the standard operating protocols. The tissue cores were then scored for immunostaining intensity. Representative images of samples were recorded by the microscope.

### TNBC orthotopic implantation mice experiment[26]

Athymic nude mice (female, aged 10 months) were obtained from the animal centre of Nanchang University and kept in a barrier facility and fed with autoclaved laboratory rodent diet. All animal studies were conducted under the principles and procedures outlined in the National regulations. MDA-MB-231 cells (or ORM2 knockdown cell lines, 1×10^7^ cells/mouse) were initially injected subcutaneously in the flank of nude mice. Small pieces of harvested subcutaneous tumor were then transplanted orthotopically into the mammary gland of nude mice. In the first set of experiments, 40 mice were randomly distributed into four groups (n=10) with different treatments: 1) control: treated with vehicle, 2)QUE-L: treated with 25 mg/kg, 3)QUE-M: treated with 50 mg/kg, 4)QUE-H: treated with 100 mg/kg. The QUE was administered by oral gavage every 3 days after orthotopic implantation. The mice were fed and treated until the end of their survival, and the survival times were recorded. In the second set of experiments, 20 mice were randomly distributed into two groups (n=10) with different treatments: 1) control: treated with vehicle, 2)QUE: treated with the best dose based on the first set of experiments. The QUE was administered by oral gavage every 3 days after orthotopic implantation. The mice were fed and treated until day 30. The body weights were recorded every 3 days and the tumor volumes were measured on day 30. Tumor samples were collected for mRNA and protein detection.

### RNA-sequencing data and bioinformatics analysis

The RNA-seq data of TCGA[27] with toil processed uniformly were downloaded from UCSC XENA[28]. This study was in full compliance with the published guidelines of TCGA. Differentially expressed gene analysis was implemented by the DESeq2 R package, comparing expression data of low- and high-expression of ORM2 (cut-off value of 50%) in TNBC samples to identify DEGs. Genes with the threshold for |logFC| >1.5 and p <0.05 were applied for functional enrichment analysis. Gene Ontology (GO) molecular functional analyses were implemented using the ClusteProfiler package in R. The protein-protein interaction networks were constructed using STRING[29].

### Statistics

Statistical analysis of all data was carried out by a two-tailed Student’s t-test. Differences were considered statistically significant if the P-value was less than 0.05.

## Results

### QUE inhibited the viability of breast cancer cell lines

In this study, we used the MTT assay, a commonly used cell viability assay[30] to measure the IC50 of QUE in TNBC cell line viability. Cells were treated with various concentrations of QUE for 12 hours prior to the assay. The study included five triple-negative breast cancer (TNBC) cell lines: MDA-MB-231, MDA-MB-468, MDA-MB-436, BT-20, and BT-549. Additionally, two non-TNBC cell lines, MDA-MB-435 (HER-enriched breast cancer) and MCF-7 (ER-positive breast cancer), as well as two non-tumorigenic epithelial breast cell lines, MCF-10A and MCF-12A, were included. Results demonstrated that QUE significantly inhibited the viability of all TNBC cell lines, as well as the HER-enriched breast cancer cell line MDA-MB-435 and the ER-positive breast cancer cell line MCF-7. The IC50 values for TNBC cell lines ranged from 15.3 μM to 55.2 μM, whereas the IC50 values for the two normal breast cell lines exceeded 100 μM. Among the TNBC cell lines, MDA-MB-231 and BT-20 exhibited the highest sensitivity to QUE, with IC50 values of 15.3 μM and 20.1 μM, respectively, while BT-549 showed the lowest sensitivity, with an IC50 of 55.2 μM. To further confirm the insensitivity of normal breast cells at low concentrations, cells were exposed to QUE for 36 hours and 72 hours, revealing IC50 values still exceeding 100 μM for the normal breast cells. Thus, it was concluded that normal breast cells were not significantly affected by QUE at concentrations below 64 μM. These findings suggested that QUE exhibited relative specificity in inhibiting breast cancer cells, although the differential effect between TNBC and non-TNBC cells may not be substantial. Subsequent experiments utilized MDA-MB-231 and BT-20 (the TNBC cell lines with the highest IC50 values) as well as MCF-10A and MCF-12A (the normal control cell lines), with QUE administered at a concentration of 64 μM.

### ORM2 was associated with TNBC and worse survival in TNBC patients

ORM2 is one of the potential targets of QUE provided by the HERB database[31], a high-throughput experiment- and reference-guided database of traditional Chinese medicine. We queried “ORM2” for the drug target and the database search results came out with three drugs, including meletin, plumb, and quercitin. We then search for “quercitin” in the PubChem database and found that it is the same as “quercetin” (PubChem id: 5280343).

According to TCGA data, ORM2 was found to be significantly upregulated in breast cancer compared to normal breast tissues. Furthermore, within breast cancer subtypes, TNBC exhibited higher ORM2 expression compared to non-TNBC (Figure 2A). We further investigated the association between ORM2 levels and different pN stages in TNBC and non-TNBC. The results revealed that N2 TNBC had higher ORM2 expression compared to N1, and N1 TNBC showed higher ORM2 expression compared to N0. However, in non-TNBC, there were no significant differences observed among different pN stages (Figure 2B). These findings suggest that ORM2 may be associated with TNBC metastasis or cell survival during migration, but it may not impact the migration of non-TNBC cells.

**Figure 1.**
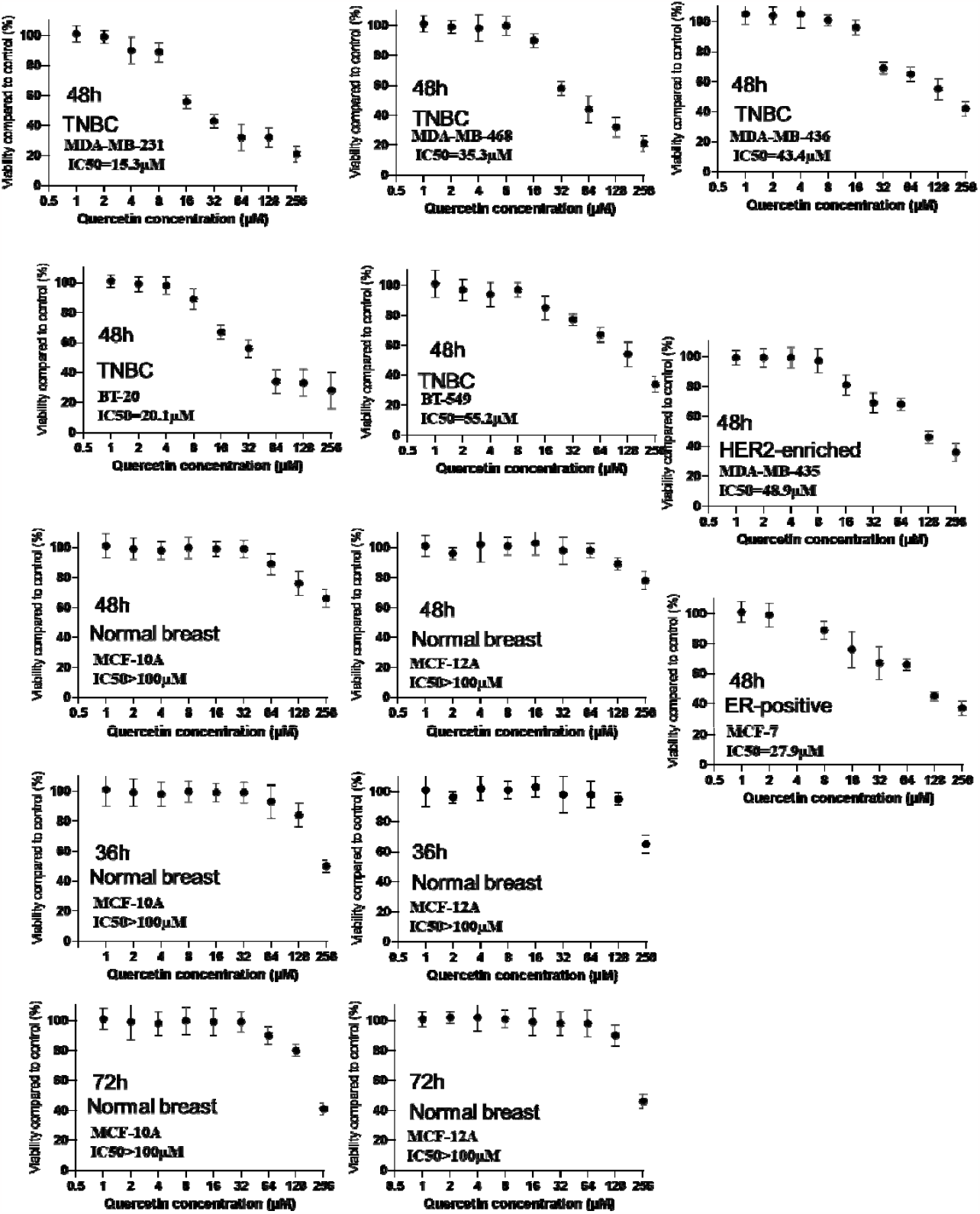
The IC50 of QUE in TNBC cell lines viability with two non-TNBC cell lines and two non-tumorigenic epithelial breast cell lines as controls.

**Figure 2.**
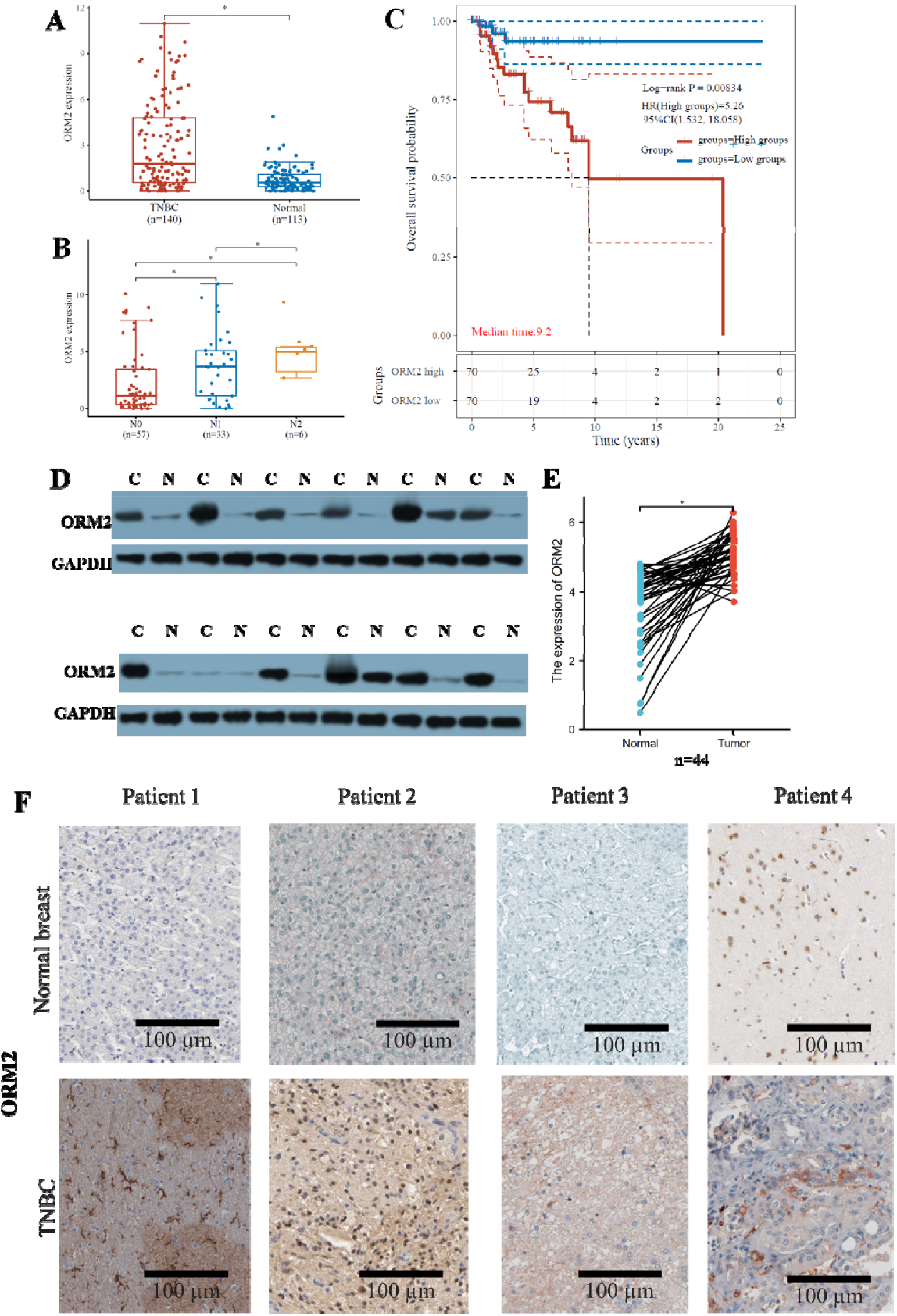
Clinical association of ORM2 and TNBC. A. Expression of ORM2 in TNBC, non-TNBC, and normal breast tissues from TCGA cohort. B. Expression of ORM2 in different N stages TNBC or non-TNBC tissues from TCGA cohort. C. Kaplan-Meier plotting of ORM2 high and low TNBC or non-TNBC patients. D-E. The protein expression of ORM2 in TNBC and paired normal breast adjacent tissues from patients. Western blotting was used to analyze the protein expression. D. Representative images of western blotting. E. Link plotting of the protein expression of ORM2 in TNBC and paired normal breast adjacent tissues. F. Representative images of immunohistochemistry stainings from TNBC and paired normal breast adjacent tissues from patients. C: cancer, N: normal. *p<0.05, ****p<0.00001

Moreover, we analyzed the overall survival of TNBC and non-TNBC patients using the TCGA cohort. The analysis demonstrated that high ORM2 expression was significantly associated with worse overall survival in TNBC but not in non-TNBC. The hazard ratio for the high ORM2 group was 5.26 in TNBC (Figure 2C). These data further support the notion that ORM2 affects TNBC but does not exert the same influence on non-TNBC. Consequently, this study focused exclusively on investigating TNBC.

To validate the overexpression of ORM2 in TNBC, we collected tumor samples and paired normal breast adjacent tissues from TNBC patients. Protein expression analysis was performed using western blotting, and protein levels were observed through immunohistochemistry staining. The results demonstrated that, overall, ORM2 protein was significantly upregulated in tumor samples compared to normal breast tissue samples. Among the 30 patients, 4 patients (13.3%) exhibited lower ORM2 protein levels in cancer tissue compared to normal tissue, while 26 patients (86.7%) showed higher ORM2 protein levels in cancer tissue (Figure 2D-E). Immunohistochemistry staining of the slides also confirmed stronger ORM2 staining in tumor samples compared to normal breast tissue samples (Figure 2F). The observed ORM2 protein overexpression was consistent with the ORM2 mRNA expression levels obtained from data mining analysis, suggesting that ORM2 protein levels are dependent on ORM2 mRNA levels. It is worth noting that the mRNA levels of ORM2 may be epigenetically regulated by transcriptional factors, requiring further exploration in future studies. These findings also indicate that TNBC may exhibit distinct ORM2 regulation compared to other breast cancer subtypes.

### ORM2 positively regulated the viability of TNBC cells but not normal breast cells

To further investigate the role of ORM2 in breast cancer, we examined ORM2 expression in seven breast cancer cell lines and two normal breast cell lines. Western blotting analysis revealed significantly higher ORM2 protein levels in the breast cancer cell lines compared to the normal breast cell lines (Figure 3A-B). Based on their high ORM2 expression, MDA-MB-231, BT-20, MCF-10A, and MCF-12A were selected for further experiments on ORM2 knockdown and overexpression. It is worth noting that both MDA-MB-231 and BT-20 are TNBC cell lines.

**Figure 3.**
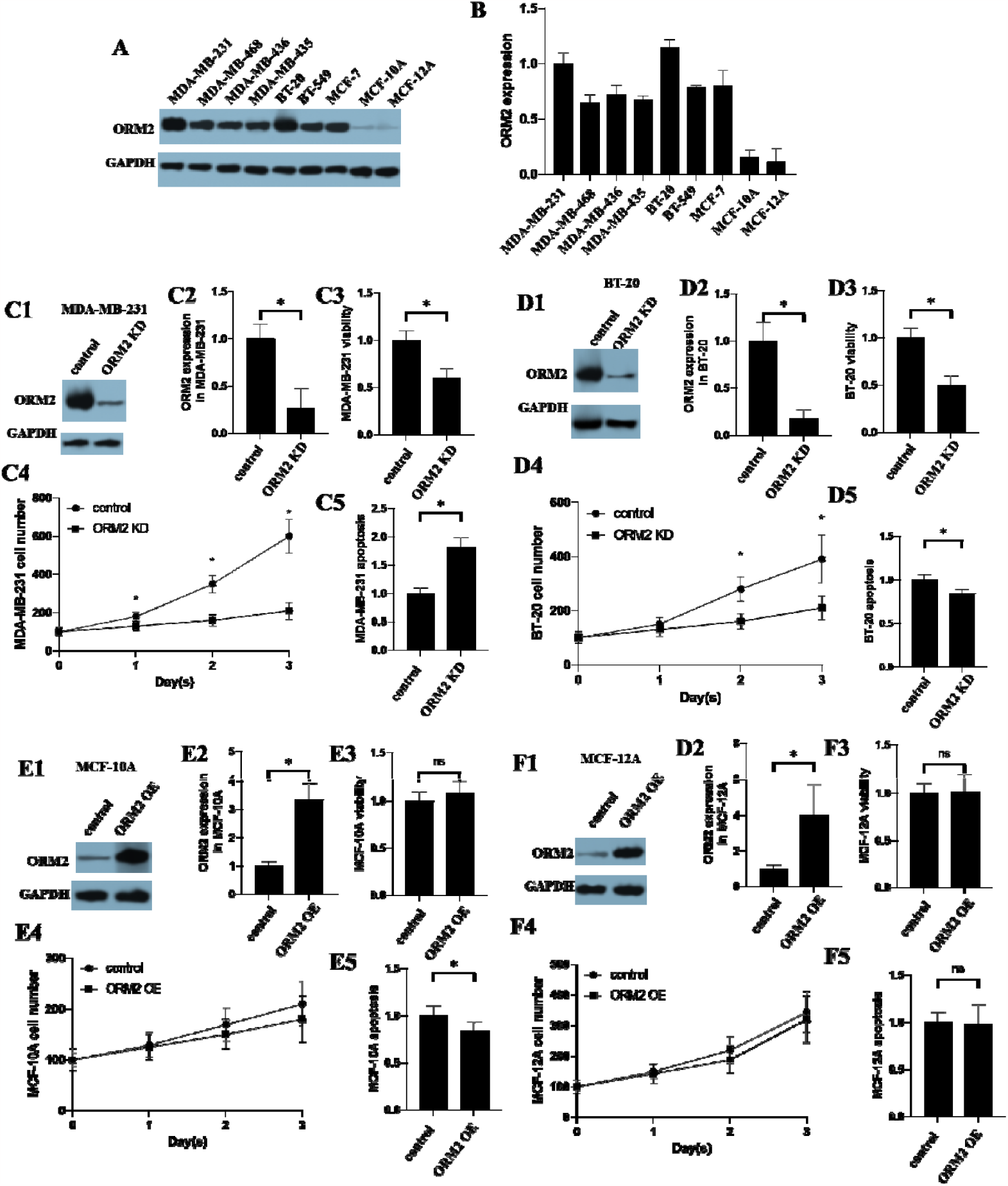
Effects of ORM2 in TNBC cell lines. A-B. The expression of ORM2 in TNBC cell lines. Two non-tumorigenic epithelial breast cell lines were used as controls. A. Representative images of western blotting. B. The bar chart of the expression of ORM2 in TNBC cell lines. B. The scatter plot and correlation of the expression of ORM2 and the IC50 of QUE in TNBC cell lines. C-F. Effects of ORM2 in ORM2-knockdown cells. C. Effects of ORM2 in MDA-MB-231. C1. Representative images of western blotting in MDA-MB-231. C2. The bar chart of the expression of ORM2 in MDA-MB-231. C3. Cell viability of MDA-MB-231. C4. Cell number counting of MDA-MB-231. C5. Apoptosis of MDA-MB-231. D. BT-20. E. MCF-10A. F. MCF-12A. D-F. legends were the same as C. *p<0.05

In this study, we successfully knocked down ORM2 expression in MDA-MB-231 and BT-20 cells, as confirmed by western blotting (Figure 3 C1-2 and D1-2). MTT assay results demonstrated that ORM2 knockdown led to reduced cell viability in both MDA-MB-231 and BT-20 cells (Figure 3 C3 and D3). Cell counting assays further confirmed a decrease in the cell number of MDA-MB-231 and BT-20 cells following ORM2 knockdown (Figure 3 C4 and D4). However, the apoptosis levels in MDA-MB-231 and BT-20 cells showed inconsistent results. ORM2 knockdown increased apoptosis in MDA-MB-231 cells but decreased apoptosis in BT-20 cells (Figure 3 C5 and D5). Considering the overall increase in cell number over time in both MDA-MB-231 and BT-20 cells, we hypothesized that the observed changes in apoptosis were not the primary factor driving the alterations in cell viability upon ORM2 knockdown. Thus, apoptosis may not play a critical role in the effect of ORM2.

Additionally, we conducted similar experiments in the two non-tumorigenic epithelial breast cell lines, MCF-10A and MCF-12A. The overexpression of ORM2 in MCF-10A and MCF-12A cells was confirmed by western blotting following vector transfection (Figure 3 E1-2 and F1-2). MTT assay results indicated that ORM2 overexpression did not affect the viability of MCF-10A and MCF-12A cells (Figure 3 E3 and F3). Cell counting assays further confirmed that ORM2 overexpression did not alter the cell number of MCF-10A and MCF-12A cells (Figure 3 E4 and F4). Furthermore, ORM2 overexpression did not significantly affect apoptosis in MCF-10A and MCF-12A cells (Figure 3 E5 and E5). These findings suggested that ORM2 positively regulated the viability of TNBC cells but did not impact normal breast cells.

### QUE inhibited the expression of ORM2 and viability in TNBC cells but not in normal breast cells

We observed the down-regulation of ORM2 protein by QUE treatment in MDA-MB-231 and BT-20 cells, as evidenced by western blotting analysis (Figure 4 A1-2 and B1-2). Cell counting assays revealed a decrease in the cell number of MDA-MB-231 and BT-20 cells following QUE treatment (Figure 4 A3 and B3), while apoptosis remained unaffected (Figure 4 A4 and B4). Conversely, QUE treatment did not affect ORM2 protein levels in MCF-10A and MCF-12A cells (Figure 4 C1-2 and D1-2). Cell counting assays demonstrated that QUE exposure did not impact the cell number of MCF-10A and MCF-12A cells (Figure 4 C4 and D4). Furthermore, QUE treatment did not significantly affect apoptosis in MCF-10A and MCF-12A cells (Figure 4 C4 and D4). These findings suggest that ORM2 positively regulates the viability of TNBC cells, while it does not exert the same effect on normal breast cells.

**Figure 4.**
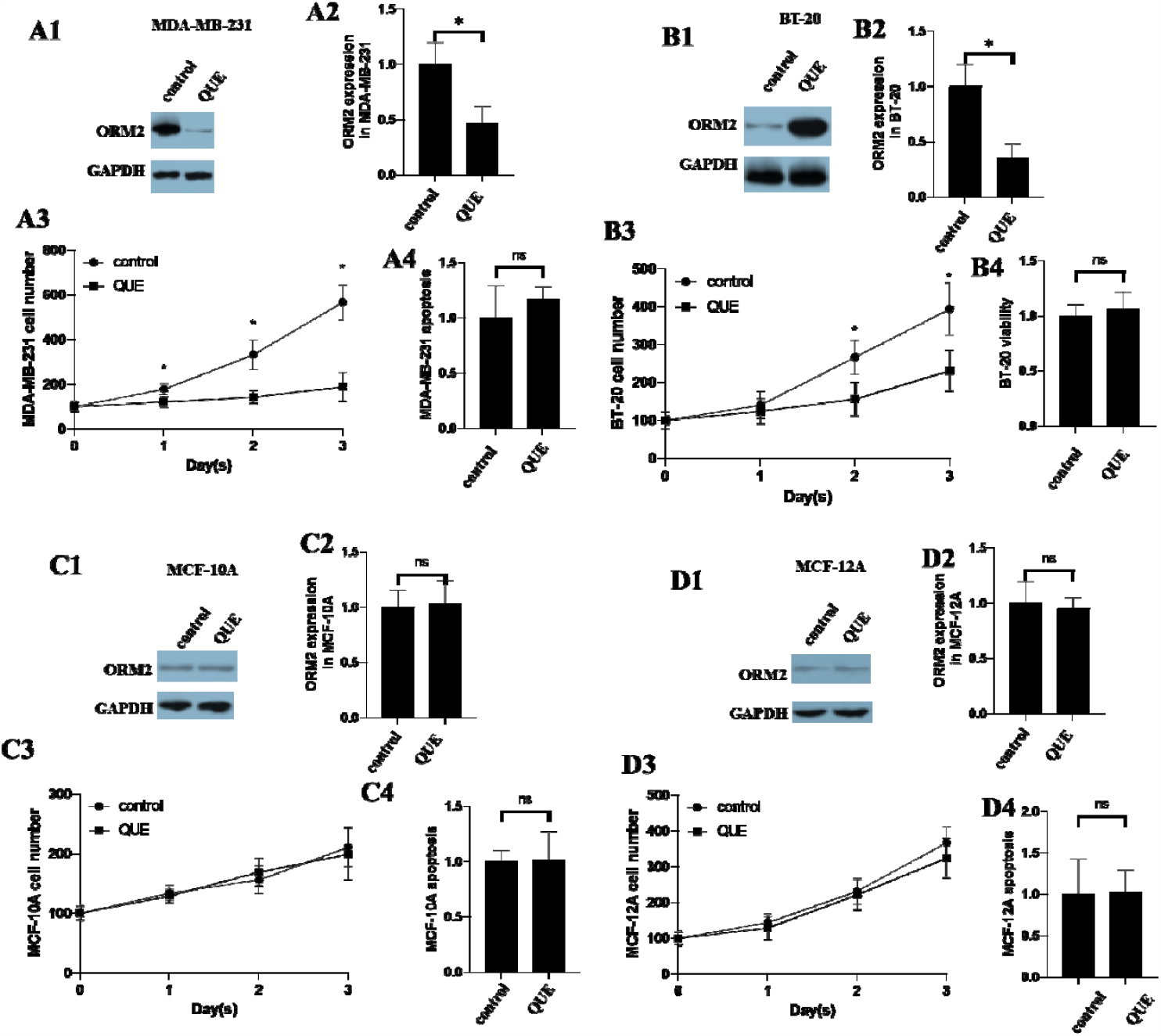
Effects of QUE in two TNBC cell lines and two non-tumorigenic epithelial breast cell lines. A. Effects of ORM2 in MDA-MB-231. A1. Representative images of western blotting in MDA-MB-231. A2. The bar chart of the expression of ORM2 in MDA-MB-231. A3. Cell number counting of MDA-MB-231. A4. Apoptosis of MDA-MB-231. B. BT-20. C. MCF-10A. D. MCF-12A. B-D. legends were the same as A. *p<0.05

### ORM2 mediated the inhibition of QUE toward the viability of TNBC cells but not normal breast cells

Correlation analysis revealed a negative correlation between ORM2 protein expression and the IC50 of QUE in breast cancer cell lines (Figure 5A). This correlation suggests that ORM2 may mediate the effect of QUE on cell viability. To validate this hypothesis in TNBC, we performed ORM2 knockdown experiments in MDA-MB-231 and BT-20 cells using two distinct siRNA constructs (KD1 and KD2). We then measured the IC50 of QUE inhibition on the viability of these cell lines. The vehicle was used as a control to account for any potential impacts of plasmid transfection on cell behavior. As depicted in Figure 5B, the IC50 of the knockdown control in MDA-MB-231 cells was 13.2 μM, whereas the IC50 of both ORM2-knockdown MDA-MB-231 cells exceeded 100 μM. Similarly, in the case of BT-20 cells, the IC50 of the knockdown control was 23.1 μM, while the IC50 of both ORM2-knockdown BT-20 cells also surpassed 100 μM. These results support the notion that ORM2 mediates the inhibitory effect of QUE on the viability of TNBC cells. Additionally, we conducted ORM2 overexpression experiments in MCF-10A and MCF-12A cells and determined the IC50 of QUE. As shown in Figure 5B, the IC50 of the knockdown control in MCF-10A cells exceeded 100 μM, and the IC50 of ORM2-knockdown MCF-10A cells was also over 100 μM. Similarly, both the IC50 of the knockdown control and ORM2-knockdown MCF-12A cells exceeded 100 μM. Even after 72 hours of QUE exposure, no significant effect on cell viability was observed in MCF-10A and MCF-12A cells with ORM2 overexpression.

**Figure 5.**
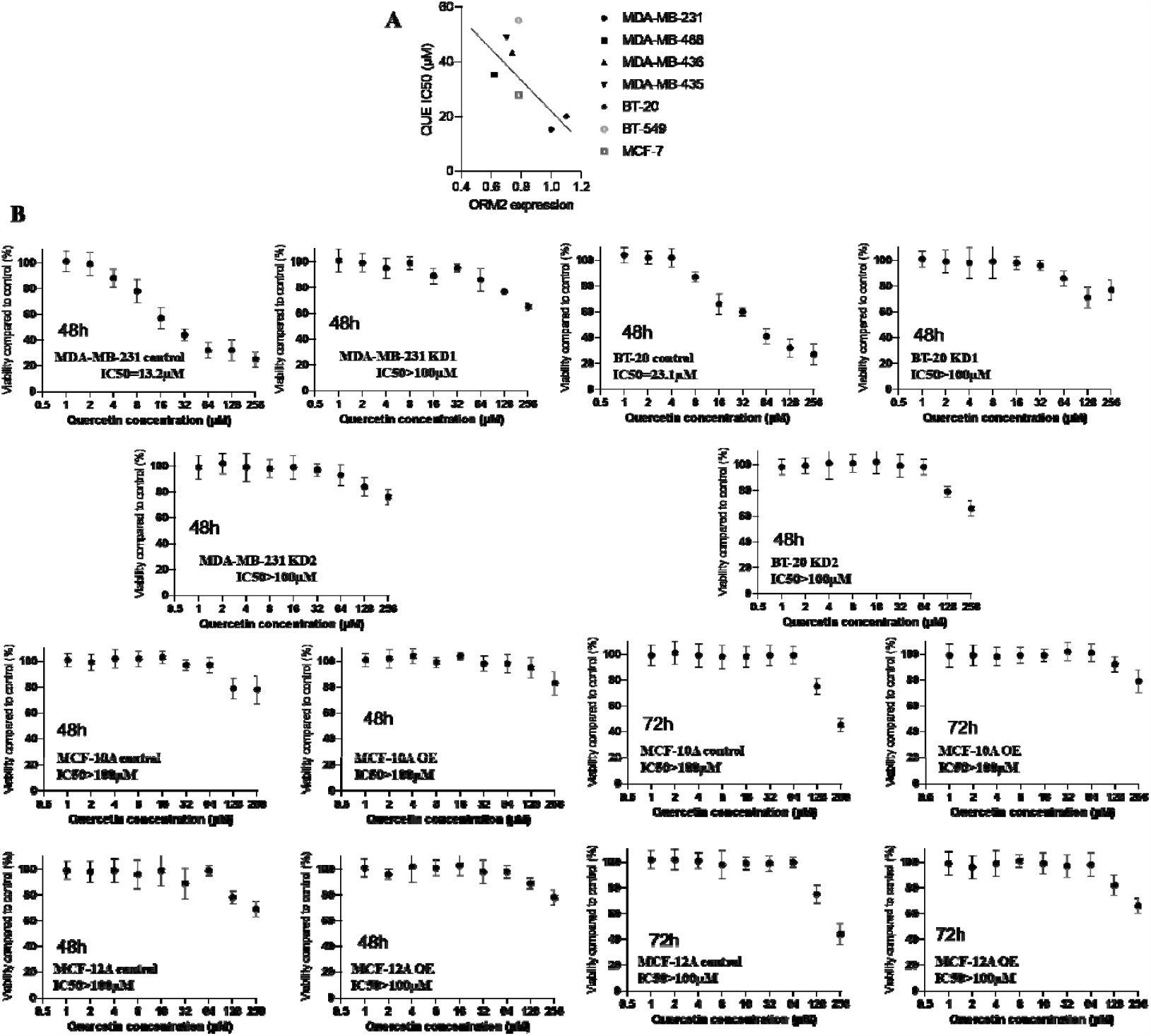
The IC50 of QUE in the viability of two ORM2-knockdown TNBC cell lines and two non-tumorigenic epithelial ORM2-overexpressing breast cell lines. MTT assay was used to determine the viability of cells. Data were normalized to the vehicle control of each cell line. A. The correlation between ORM2 expression and QUE IC50. B. IC50 of cell lines.

### QUE increased survival and decreased expression of ORM2 in tumors in TNBC mice

To evaluate the therapeutic effect of QUE on TNBC, we utilized an orthotopic implantation TNBC mouse model. Initially, we conducted a survival analysis to determine the optimal treatment dose. The QUE-L groups showed minimal effect on survival compared to the control group. However, both QUE-M and QUE-H groups significantly improved the survival of TNBC mice, with the QUE-H group demonstrating the most favorable outcomes (Figure 6A). Thus, we selected the QUE-H concentration for subsequent studies. To minimize survival bias, we chose day 30 as the endpoint when the majority of mice were still alive. We monitored the body weight of the mice every three days and measured tumor volume at the endpoint. The results indicated that the body weights of the mice did not exhibit significant changes. Although the QUE-treated groups showed slightly lower body weights compared to the control group, the difference was not statistically significant (Figure 6B). However, the tumor volume in the QUE-treated groups was significantly reduced compared to the control group (Figure 6C). Notably, the expression of ORM2 protein in the tumors was markedly decreased in the QUE-treated groups compared to the control group (Figure 6D-E). Representative images of immunohistochemistry staining from tumors in the mice were presented in Figure 6F.

**Figure 6.**
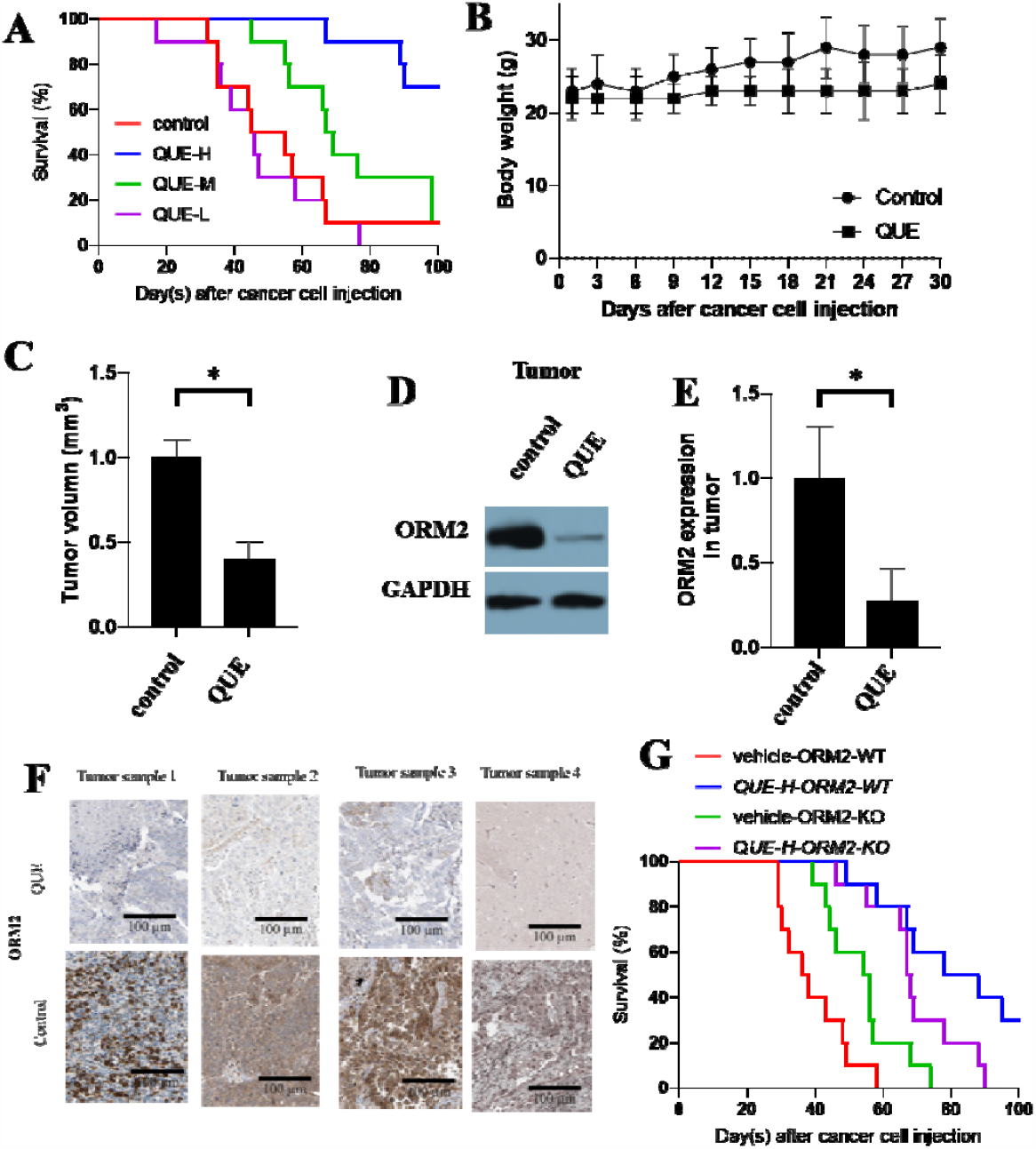
Effect of QUE on TNBC in the mouse model. A. Survival of mice with TNBC with QUE treatment. B. Body weight of mice. C. Tumor volume of mice. D-E. The expression ORM2 in tumors. F. Representative images of immunohistochemistry stainings from tumors from mice. G. The survival of ORM2 knowdown cells animals and ORM2 wild-type animals with QUE (QUE-H) treatment. *p<0.05

To investigate whether attenuating ORM2 activity influences the therapeutic effect of QUE in TNBC treatment in vivo, we employed the ORM2 knockdown cell line in the in vivo study. We compared the overall treatment effect (survival) of QUE between ORM2 knockdown cell animals and ORM2 wild-type animals. The results revealed a striking difference in treatment outcomes between the treatment and vehicle groups in the wild-type cell model. However, in the ORM2 knockdown group, the difference between the treatment and vehicle groups was less pronounced. It is important to note that this experiment was limited by the transfection duration of the cell lines. While the cells developed tumors, the ORM2 knockdown may have been rapidly recovered to the same level, thereby potentially accounting for the difference in treatment effect observed in the initial few days and the final results. Notably, the vehicle group in the ORM2 knockdown model exhibited similar survival rates as the QUE treatment group in the ORM2 wild-type model, particularly before day 50. These findings suggest that ORM2 knockdown has a comparable effect to QUE treatment, consistent with previous results indicating that QUE reduces the level of ORM2.

### Potential mechanism of QUE on ORM2

To gain insights into the potential mechanisms underlying the effect of QUE on ORM2 function in TNBC, we conducted an enrichment analysis of the differentially expressed genes associated with high and low ORM2 expression in TNBC, based on the TCGA cohort. The analysis revealed 244 up-regulated genes and 200 down-regulated genes that were associated with ORM2 expression (Figure 7A). The up-regulated genes were found to be enriched in functions such as transmembrane receptor protein kinase activity, extracellular matrix structural constituent, and growth factor binding. Conversely, the down-regulated genes were associated with functions including interleukin-15 receptor activity, CXCR3 chemokine receptor binding, TAP binding, deoxycytidine deaminase activity, and peptide antigen binding (Figure 7B).

**Figure 7.**
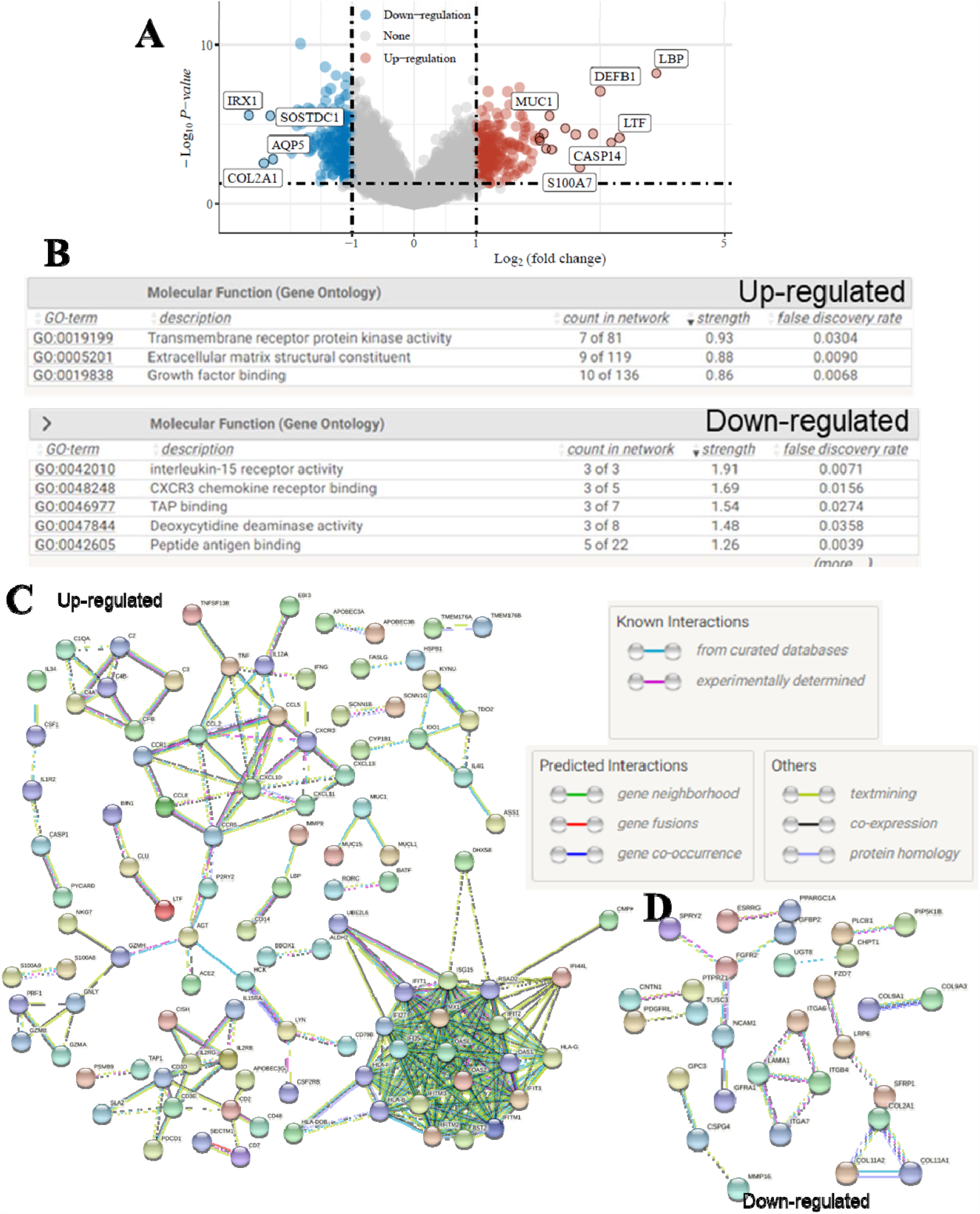
Bioinformatics analysis of ORM2 and QUE. A. Differentially expressed gene analysis of ORM2 high and low TNBC from TCGA cohort. Down-regulate: negatively associated with ORM2; up-regulate: positively associated with ORM2. B. GO enrichment analysis of Differentially expressed genes. C. Protein-protein interaction network of up-regulated differentially expressed genes. D. Protein-protein interaction network of down-regulated differentially expressed genes.

Based on these differentially expressed genes, we constructed protein-protein interaction (PPI) networks. The PPI network for the up-regulated genes consisted of 242 nodes and 270 edges, with an average node degree of 2.23 and an average local clustering coefficient of 0.325. The PPI network for the down-regulated genes comprised 197 nodes and 22 edges, with an average node degree of 0.223 and an average local clustering coefficient of 0.129. The PPI enrichment analysis indicated a highly significant p-value of < 1.0e-16 for the up-regulated genes network and a significant p-value of < 9.22e-06 for the down-regulated genes network (Figure 7C-D). These findings provide a comprehensive view of the potential interactions and functional associations of genes differentially expressed in relation to ORM2 in TNBC.

## Discussions

Traditional medicine has been suggested to be a potential resource to discover therapeutic drugs for human diseases [32-38]. Traditional medicine is widely used as an alternative treatment for cancer and is especially popular in Asian countries such as China, Japan, and Korea [39]. Traditional medicine can have multiple effects on cancers [40] and have the unique advantages of multi targets and fewer side effects, therefore, it is thought to be safe for long-term complementary and alternative therapies for TNBC [41]. QUE is one of the promising drug candidates for cancer treatment from traditional medicine. QUE is the major representative of the flavonoid subclass of flavonols and is ubiquitously present in daily fruits and vegetables, such as onions. According to the data from the US Department of Health and Human Services, QUE is one of the most common dietary flavonols in the US with an average daily intake of approximately 25 mg [42]. Drug dose is one of the critical issues in clinical treatment [30, 43, 44]. In our animal experiments, we determined that the lowest effective dose of QUE was 50 mg/kg (QUE-M), which was only twice the average daily intake amount. Although mice and humans differ, these data suggest that the effective concentration range of QUE might be within an acceptable range for normal intake. The recorded body weight of the mice showed no significant reduction, even at a dose of 100 mg/kg (QUE-H), indicating that the QUE treatment might not be toxic to the mice. Furthermore, the IC50 values of the two normal breast cell lines were over 100 μM, which was much higher than the IC50 values observed for TNBC cell lines. These results indicate that cancer cells are much more sensitive to QUE than normal cells. This finding aligns with our animal study, where we concluded that QUE specifically targets TNBC cells and is not toxic to the animals.

ORM2 has been suggested to associate with colon cancer[45, 46]. The plasma ORM2 levels were reported to be used for the prognosis of colorectal cancer patients[45], which indicated that the ORM2 was presented in plasma. The development of colon cancer is associated with cancer stem cells[47], thus, whether ORM2 is associated with cancer stemness can be investigated in the future. Additionally, a previous study revealed that the downregulation of ORM2 acted as a prognostic factor associated with cancer-promoting pathways in liver cancer[48]. Our results that ORM2 in breast cancer was much higher than that in normal tissues and the results that ORM2 was associated with the poor survival rate strongly supported the potential diagnostic and prognostic values of ORM2 for TNBC patients. As blood is a much more accessible clinical sample than tissues, the investigation of the plasma ORM2 levels in TNBC patients would be conducive to the development of ORM2 as a diagnostic and prognostic biomarker of TNBC. Bioinformatic analysis has been widely used to evaluate the prognostic value of genes[49-55], in this study, our bioinformatic analysis revealed that ORM2 was a potential prognostic biomarker for TNBC but not for non-TNBC.

In this study, we did not have kinetic evidence to determine the turnover of ORM2 proteins in cells, which prevented us from identifying the exact mechanism by which QUE influences the expression or degradation of ORM2 protein. However, our findings that QUE influences the level of ORM2 protein and that ORM2 mediates the development of TNBC are still of significant scientific importance. While the specific mechanism of ORM2 regulation remains unclear, we conducted differential expressed gene analysis, enrichment analysis, and constructed a protein-protein interaction network to gain insights into potential regulatory mechanisms. The associations we identified might shed light on how ORM2 is regulated or how ORM2 regulates other proteins. It is important to note that the specific function of ORM2 is still not fully understood at this stage. A study has shown that ORM2 appears to play a role in regulating the activity of the immune system during the acute-phase reaction [56], which might function in the early development of cancer. Accumulating data suggested that cancer-associated inflammation can facilitate cancer development and progression [43, 57, 58]. In turn, cancer could induce cancer-associated inflammation, which may play a pivotal role in all stages of tumorigenesis [59, 60]. A novel finding of this study was that ORM2 might play an extra role in cancer cells besides cancer-associated inflammation. In our in vitro study, no immune cells were presented but the viability of cells remained affected by ORM2. The viability of cancer cells can be affected by apoptosis, proliferation, necrosis, autophagy, etc. It is not sure which aspects of the cell viability were affected by ORM2. In this study, we have tested apoptosis. We suggested that the effect of ORM2 on cell viability was not mediated by apoptosis. Exploring the underlying mechanism of ORM2 effects on cancer cell viability is required. Although this study did not go deeply into this topic, the enrichment analysis and PPI network might provide some valuable hints for future studies as some previous studies did [49, 50, 55, 61].

The molecular biological technique has been used wildly to manipulate the expression of a target protein[62]. Our knockdown experiments revealed that the effect of QUE on TNBC cell viability was influenced by the reduction of ORM2. On the other hand, our results also showed that this mechanism might not be the case in normal breast cells. These results were consistent with the previous assumption that QUE is specific to TNBC cells but not normal breast cells. This is because the ORM2 viability regulation might be present only in TNBC cells but not in normal breast cells. This evidence supported ORM2 as an effective and specific target of QUE in TNBC treatment. In addition, drugs used in clinical cancer treatment can have potential effects on immunity[63-65]. Mice in this study were immune definition mice, which also supported the extra effect of ORM2 besides immunity regulations. Nevertheless, laboratory evidence is required to conclude the potential mechanism of QUE on ORM2, therefore, we carefully state that the pathways identified were “potential mechanisms”. We hope we did not overstate our analysis results and mislead the readers. In the bioinformatics analysis, the associations between ORM2 and other proteins are mainly based on co-presenting or negative correlation. Therefore the causal effect of ORM2 on these pathways was not clear at all. Further studies are required to obtain laboratory evidence of these potential mechanisms. Interestingly, GETx data suggested that ORM2 is highly expressed in the liver (S-Figure). Previous studies have revealed that QUE might protect the liver from injury[66-69]. Given the fact found in this study that ORM2 is a direct target of QUE, it is not sure if ORM2 play a role in the impact of QUE on the liver. We proposed that this is an interesting scientific question that should be explored in the future.

To sum up, QUE inhibited cell viability via ORM2 in TNBC. Our study is conducive to a better understanding of QUE effects on TNBC and contributes to the further development of QUE treatment for TNBC patients.

## Ethical approval

Ethical approval was sought from an ethics committee of the Second Affiliated Hospital of Nanchang University.

## Acknowledgements

The authors would like to express their gratitude to the Second Affiliated Hospital of Nanchang University and the Jiangxi Tumor Hospital.

## Authors’ contributions

Aiqing Zhou and Zhijun Chen designed the study. Yan Wang conducted the experiments. Liling Tan and Long Chen wrote the original manuscript. Lin Wang downloaded and analyzed the data. Zhijun Chen supervised the study and edited the manuscript.

## Availability of data and materials

The datasets analyzed during the present study are available from the corresponding author Zhijun Chen.

## References

[1] Bray F, Laversanne M, Weiderpass E and Soerjomataram I. The ever-increasing importance of cancer as a leading cause of premature death worldwide. Cancer 2021; 127: 3029–3030.

[2] Jemal A, Bray F, Center MM, Ferlay J, Ward E and Forman D. Global cancer statistics. CA Cancer J Clin 2011; 61: 69–90.

[3] Siegel RL, Miller KD, Fuchs HE and Jemal A. Cancer Statistics, 2021. CA Cancer J Clin 2021; 71: 7–33.

[4] Liu H, Dilger JP and Lin J. The Role of Transient Receptor Potential Melastatin 7 (TRPM7) in Cell Viability: A Potential Target to Suppress Breast Cancer Cell Cycle. Cancers (Basel) 2020; 12:

[5] Vagia E, Mahalingam D and Cristofanilli M. The Landscape of Targeted Therapies in TNBC. Cancers (Basel) 2020; 12:

[6] Liu H. Nav channels in cancers: Nonclassical roles. Global Journal of Cancer Therapy 2020; 6: 5.

[7] Subramani R, Nandy SB, Pedroza DA and Lakshmanaswamy R. Role of Growth Hormone in Breast Cancer. Endocrinology 2017; 158: 1543–1555.

[8] Scher HI, Steineck G and Kelly WK. Hormone-refractory (D3) prostate cancer: refining the concept. Urology 1995; 46: 142–148.

[9] Keegan TH, Kurian AW, Gali K, Tao L, Lichtensztajn DY, Hershman DL, Habel LA, Caan BJ and Gomez SL. Racial/ethnic and socioeconomic differences in short-term breast cancer survival among women in an integrated health system. Am J Public Health 2015; 105: 938–946.

[10] Keegan TH, Kurian AW, Gali K, Tao L, Lichtensztajn DY, Hershman DL, Habel LA, Caan BJ and Gomez SL. Racial/ethnic and socioeconomic differences in short-term breast cancer survival among women in an integrated health system. American journal of public health 2015; 105: 938–946.

[11] Liu H, Dilger JP and Lin J. Lidocaine Suppresses Viability and Migration of Human Breast Cancer Cells: TRPM7 as A Target for Some Breast Cancer Cell Lines. Cancers (Basel) 2021; 13: 234.

[12] Liu H. A prospective for the role of two-pore channels in breast cancer cells. Global Journal of Cancer Therapy 2020; 6: 001--003.

[13] Swain SM, Kim S-B, Cortés J, Ro J, Semiglazov V, Campone M, Ciruelos E, Ferrero J-M, Schneeweiss A and Knott A. Pertuzumab, trastuzumab, and docetaxel for HER2-positive metastatic breast cancer (CLEOPATRA study): overall survival results from a randomised, double-blind, placebo-controlled, phase 3 study. The lancet oncology 2013; 14: 461–471.

[14] Suolinna E, Buchsbaum R and Racker E. The effect of flavonoids on aerobic glycolysis and growth of tumor cells. Cancer Research 1975; 35: 1865–1872.

[15] Ezzati M, Yousefi B, Velaei K and Safa A. A review on anti-cancer properties of Quercetin in breast cancer. Life Sci 2020; 248: 117463.

[16] Kundur S, Prayag A, Selvakumar P, Nguyen H, McKee L, Cruz C, Srinivasan A, Shoyele S and Lakshmikuttyamma A. Synergistic anticancer action of quercetin and curcumin against triple-negative breast cancer cell lines. J Cell Physiol 2019; 234: 11103–11118.

[17] Umar SM, Patra S, Kashyap A, Dev JRA, Kumar L and Prasad CP. Quercetin Impairs HuR-Driven Progression and Migration of Triple Negative Breast Cancer (TNBC) Cells. Nutr Cancer 2021; 1-14.

[18] Srinivasan A, Thangavel C, Liu Y, Shoyele S, Den RB, Selvakumar P and Lakshmikuttyamma A. Quercetin regulates β-catenin signaling and reduces the migration of triple negative breast cancer. Mol Carcinog 2016; 55: 743–756.

[19] Luo Z, Lei H, Sun Y, Liu X and Su DF. Orosomucoid, an acute response protein with multiple modulating activities. J Physiol Biochem 2015; 71: 329–340.

[20] Dalby B, Cates S, Harris A, Ohki EC, Tilkins ML, Price PJ and Ciccarone VC. Advanced transfection with Lipofectamine 2000 reagent: primary neurons, siRNA, and high-throughput applications. Methods 2004; 33: 95–103.

[21] Li R, Xiao C, Liu H, Huang Y, Dilger JP and Lin J. Effects of local anesthetics on breast cancer cell viability and migration. BMC cancer 2018; 18: 666.

[22] Salgame P, Varadhachary AS, Primiano LL, Fincke JE, Muller S and Monestier M. An ELISA for detection of apoptosis. Nucleic acids research 1997; 25: 680–681.

[23] Crowley LC, Marfell BJ, Christensen ME and Waterhouse NJ. Measuring Cell Death by Trypan Blue Uptake and Light Microscopy. Cold Spring Harb Protoc 2016; 2016:

[24] Liu X, Liu H, Xiong Y, Yang L, Wang C, Zhang R and Zhu X. Postmenopausal osteoporosis is associated with the regulation of SP, CGRP, VIP, and NPY. Biomed Pharmacother 2018; 104: 742–750.

[25] Gusti-Ngurah-Putu EP, Huang L and Hsu YC. Effective Combined Photodynamic Therapy with Lipid Platinum Chloride Nanoparticles Therapies of Oral Squamous Carcinoma Tumor Inhibition. J Clin Med 2019; 8:

[26] Sommaggio R, Cappuzzello E, Dalla Pietà A, Tosi A, Palmerini P, Carpanese D, Nicolè L and Rosato A. Adoptive cell therapy of triple negative breast cancer with redirected cytokine-induced killer cells. Oncoimmunology 2020; 9: 1777046.

[27] Tomczak K, Czerwińska P and Wiznerowicz M. The Cancer Genome Atlas (TCGA): an immeasurable source of knowledge. Contemp Oncol (Pozn) 2015; 19: A68–77.

[28] Wang S, Xiong Y, Zhao L, Gu K, Li Y, Zhao F, Li J, Wang M, Wang H, Tao Z, Wu T, Zheng Y, Li X and Liu XS. UCSCXenaShiny: An R/CRAN Package for Interactive Analysis of UCSC Xena Data. Bioinformatics 2021; 38: 527–529.

[29] von Mering C, Jensen LJ, Snel B, Hooper SD, Krupp M, Foglierini M, Jouffre N, Huynen MA and Bork P. STRING: known and predicted protein-protein associations, integrated and transferred across organisms. Nucleic Acids Res 2005; 33: D433–437.

[30] Liu H, Dilger JP and Lin J. Effects of local anesthetics on cancer cells. Pharmacology & Therapeutics 2020; 212: 107558.

[31] Fang S, Dong L, Liu L, Guo J, Zhao L, Zhang J, Bu D, Liu X, Huo P, Cao W, Dong Q, Wu J, Zeng X, Wu Y and Zhao Y. HERB: a high-throughput experiment- and reference-guided database of traditional Chinese medicine. Nucleic Acids Res 2021; 49: D1197–d1206.

[32] Haixia W, Shu M, Li Y, Panpan W, Kehuan S, Yingquan X, Hengrui L, Xiaoguang L, Zhidi W and Ling O. Effectiveness associated with different therapies for senile osteopo-rosis: a network Meta-analysis. J Tradit Chin Med 2020; 40: 17–27.

[33] Liu H, Xiong Y, Zhu X, Gao H, Yin S, Wang J, Chen G, Wang C, Xiang L, Wang P, Fang J, Zhang R and Yang L. Icariin improves osteoporosis, inhibits the expression of PPARgamma, C/EBPalpha, FABP4 mRNA, N1ICD and jagged1 proteins, and increases Notch2 mRNA in ovariectomized rats. Exp Ther Med 2017; 13: 1360–1368.

[34] Chen G, Wang C, Wang J, Yin S, Gao H, Xiang LU, Liu H, Xiong Y, Wang P, Zhu X, Yang LI and Zhang R. Antiosteoporotic effect of icariin in ovariectomized rats is mediated via the Wnt/beta-catenin pathway. Exp Ther Med 2016; 12: 279–287.

[35] Wang C, Chen G, Wang J, Liu H, Xiong Y, Wang P, Yang L, Zhu X and Zhang R. Effect of Herba Epimedium Extract on Bone Mineral Density and Microstructure in Ovariectomised Rat. Journal of Pharmaceutical and Biomedical Sciences 2016; 6:

[36] Wu Z, Ou L, Wang C, Yang L, Wang P, Liu H, Xiong Y, Sun K, Zhang R and Zhu X. Icaritin induces MC3T3-E1 subclone14 cell differentiation through estrogen receptor-mediated ERK1/2 and p38 signaling activation. Biomed Pharmacother 2017; 94: 1–9.

[37] Liu H, Xiong Y, Wang H, Yang L, Wang C, Liu X, Wu Z, Li X, Ou L, Zhang R and Zhu X. Effects of water extract from epimedium on neuropeptide signaling in an ovariectomized osteoporosis rat model. J Ethnopharmacol 2018; 221: 126–136.

[38] Liu H. Toxic medicine used in Traditional Chinese Medicine for cancer treatment: are ion channels involved? Journal of Traditional Chinese Medicine 2022; 42: 1019–1022.

[39] Xiang Y, Guo Z, Zhu P, Chen J and Huang Y. Traditional Chinese medicine as a cancer treatment: Modern perspectives of ancient but advanced science. Cancer Med 2019; 8: 1958–1975.

[40] Liu H. Effect of Traditional Medicine on Clinical Cancer. Biomedical Journal of Scientific & Technical Research 2020; 30: 23548–23551.

[41] Yang Z, Zhang Q, Yu L, Zhu J, Cao Y and Gao X. The signaling pathways and targets of traditional Chinese medicine and natural medicine in triple-negative breast cancer. J Ethnopharmacol 2021; 264: 113249.

[42] Stavric B. Quercetin in our diet: from potent mutagen to probable anticarcinogen. Clinical biochemistry 1994; 27: 245–248.

[43] Jin Z, Zhang W, Liu H, Ding A, Lin Y, Wu SX and Lin J. Potential Therapeutic Application of Local Anesthetics in Cancer Treatment. Recent Pat Anticancer Drug Discov 2022;

[45] Gao F, Zhang X, Whang S and Zheng C. Prognostic impact of plasma ORM2 levels in patients with stage II colorectal cancer. Ann Clin Lab Sci 2014; 44: 388–393.

[46] Zhang X, Xiao Z, Liu X, Du L, Wang L, Wang S, Zheng N, Zheng G, Li W, Zhang X, Dong Z, Zhuang X and Wang C. The potential role of ORM2 in the development of colorectal cancer. PLoS One 2012; 7: e31868.

[47] Liu H. A Prospective for the Potential Effect of Local Anesthetics on Stem-Like Cells in Colon Cancer. Biomedical Journal of Scientific & Technical Research 2020; 25: 18927–18930.

[48] Zhu HZ, Zhou WJ, Wan YF, Ge K, Lu J and Jia CK. Downregulation of orosomucoid 2 acts as a prognostic factor associated with cancer-promoting pathways in liver cancer. World J Gastroenterol 2020; 26: 804–817.

[49] Liu H and Weng J. A Pan-Cancer Bioinformatic Analysis of RAD51 Regarding the Values for Diagnosis, Prognosis, and Therapeutic Prediction. Frontiers in Oncology 2022; 12:

[50] Liu H and Weng J. A Comprehensive Bioinformatic Analysis of Cyclin-dependent Kinase 2 (CDK2) in Glioma. Gene 2022; 146325.

[51] Liu H and Tang T. Pan-cancer genetic analysis of cuproptosis and copper metabolism-related gene set. Front Oncol 2022; 12: 952290.

[52] Liu H and Li Y. Potential roles of Cornichon Family AMPA Receptor Auxiliary Protein 4 (CNIH4) in head and neck squamous cell carcinoma. Cancer Biomark 2022;

[53] Liu H, Dilger JP and Lin J. A pan-cancer-bioinformatic-based literature review of TRPM7 in cancers. Pharmacology & Therapeutics 2022; 108302.

[54] Liu H. Pan-cancer profiles of the cuproptosis gene set. Am J Cancer Res 2022; 12: 4074–4081.

[55] Li Y and Liu H. Clinical powers of Aminoacyl tRNA Synthetase Complex Interacting Multifunctional Protein 1 (AIMP1) for head-neck squamous cell carcinoma. Cancer Biomark 2022;

[56] Wan JJ, Wang PY, Zhang Y, Qin Z, Sun Y, Hu BH, Su DF, Xu DP and Liu X. Role of acute-phase protein ORM in a mice model of ischemic stroke. J Cell Physiol 2019; 234: 20533–20545.

[57] Mantovani A, Allavena P, Sica A and Balkwill F. Cancer-related inflammation. Nature 2008; 454: 436–444.

[58] Singh N, Baby D, Rajguru JP, Patil PB, Thakkannavar SS and Pujari VB. Inflammation and cancer. Ann Afr Med 2019; 18: 121–126.

[59] Coussens LM and Werb Z. Inflammation and cancer. Nature 2002; 420: 860–867.

[60] Diakos CI, Charles KA, McMillan DC and Clarke SJ. Cancer-related inflammation and treatment effectiveness. Lancet Oncol 2014; 15: e493–503.

[61] Li Y, Liu H and Han Y. Potential Roles of Cornichon Family AMPA Receptor Auxiliary Protein 4 (CNIH4) in Head and Neck Squamous Cell Carcinoma. Research Square 2021;

[62] Li X, Peng B, Zhu X, Wang P, Xiong Y, Liu H, Sun K, Wang H, Ou L, Wu Z, Liu X, He H, Mo S, Peng X, Tian Y, Zhang R and Yang L. Changes in related circular RNAs following ERbeta knockdown and the relationship to rBMSC osteogenesis. Biochem Biophys Res Commun 2017; 493: 100–107.

[64] Liu H. A clinical mini-review: Clinical use of Local anesthetics in cancer surgeries. The Gazette of Medical Sciences 2020; 1: 030–034.

[65] Li R, Liu H, Dilger JP and Lin J. Effect of Propofol on breast Cancer cell, the immune system, and patient outcome. BMC Anesthesiol 2018; 18: 77.

[66] Chen X. Protective effects of quercetin on liver injury induced by ethanol. Pharmacognosy magazine 2010; 6: 135.

[67] Pavanato A, Tuñón MJ, Sánchez-Campos S, Marroni CA, Llesuy S, González-Gallego J and Marroni N. Effects of quercetin on liver damage in rats with carbon tetrachloride-induced cirrhosis. Digestive diseases and sciences 2003; 48: 824–829.

[68] Molina MF, Sanchez-Reus I, Iglesias I and Benedi J. Quercetin, a flavonoid antioxidant, prevents and protects against ethanol-induced oxidative stress in mouse liver. Biological and Pharmaceutical Bulletin 2003; 26: 1398–1402.

[69] Zhao X, Wang J, Deng Y, Liao L, Zhou M, Peng C and Li Y. Quercetin as a protective agent for liver diseases: A comprehensive descriptive review of the molecular mechanism. Phytotherapy Research 2021; 35: 4727–4747.

